# Identification of translocation inhibitors targeting the type III secretion system of enteropathogenic *Escherichia coli*

**DOI:** 10.1101/2021.05.11.443714

**Authors:** Sabrina Mühlen, Viktor A. Zapol’skii, Ursula Bilitewski, Petra Dersch

## Abstract

Infections with enteropathogenic *E. coli* (EPEC) cause severe diarrhea in children. The non-invasive bacteria adhere to enterocytes of the small intestine and use a type III secretion system (T3SS) to inject effector proteins into host cells to modify and exploit cellular processes in favor of bacterial survival and replication. Several studies have shown that the T3SSs of bacterial pathogens are essential for virulence. Furthermore, the loss of T3SS-mediated effector translocation results in increased immune recognition and clearance of the bacteria. The T3SS is, therefore, considered a promising target for antivirulence strategies and novel therapeutics development. Here, we report the results of a high-throughput screening assay based on the translocation of the EPEC effector protein Tir. Using this assay, we screened more than 13,000 small molecular compounds of six different compound libraries and identified three substances which showed a significant dose-dependent effect on translocation without adverse effects on bacterial or eukaryotic cell viability. Additionally, these substances reduced bacterial binding to host cells, effector-dependent cell detachment and abolished A/E lesion formation without affecting the expression of components of the T3SS or associated effector proteins. Moreover, no effects of the inhibitors on bacterial motility or Shiga-toxin expression were observed. In summary, we have identified three new compounds that strongly inhibit T3SS-mediated translocation of effectors into mammalian cells, which could be valuable as lead substances for treating EPEC and EHEC infections.

## INTRODUCTION

Infection with enteropathogenic *Escherichia coli* (EPEC) is a major cause of infantile diarrhea in children in developing countries (1). EPEC is a member of the family of attaching and effacing (A/E) pathogens, which also includes enterohaemorrhagic *E. coli* (EHEC) and the mouse pathogen *Citrobacter rodentium*. The bacteria in this family intimately adhere to the surface of enterocytes by inducing the formation of characteristic actin-rich pedestals (attachment) and the loss of microvilli (effacement) (2). Subversion of the host actin signaling is mediated by the injection of bacterial effector proteins via a type III secretion system (T3SS). Both, the components of the T3SS, as well as effectors involved in A/E lesion formation, are encoded on the locus of enterocyte effacement pathogenicity island (LEE) located in the bacterial genome (3, 4).

The T3SS of EPEC consists of the basal body, needle, filament and translocon pore (5). The needle is formed by multiple copies of EscF, which form a hollow tube (6) essential for T3S and thus for virulence (7). It is associated on one end with the basal body that powers the translocation and on the other end with a filament made of EspA multimers (6). Together, needle and filament form a channel that upon host cell contact connects the bacterial cytosol with the host cell membrane and initiates the formation of the translocation pore. The translocon pore is formed by heterooligomers of EspB and EspD subunits that are translocated through the channel to the top of which they bind. This complex then inserts into the host cell membrane to form the pore for effector translocation (8). In addition to the T3SS, most translocated effector proteins that are essential for intimate attachment and pathogenesis are located on the LEE. The trans-located intimin receptor (Tir) is the most important and best-studied effector protein of EPEC. After translocation, it inserts into the host cell membrane and acts as a receptor for the bacterial outer membrane protein intimin (9, 10). EPEC strains with a non-functional secretion system, such as Δ*escN* and Δ*espA* mutants, are unable to intimately adhere to enterocytes and were shown to be avirulent in infection models (7).

T3SSs are highly conserved and shared by more than 25 human, plant and zoonotic Gram-negative pathogens (11). Loss of a functional T3SS was also linked with avirulence in the human enteric pathogens *Salmonella*, *Yersinia* and *Shigella* sp. (11). High conservation among the structural components of the T3SSs (11) makes it likely that inhibitors may be found that target the T3SS of more than one bacterial pathogen. Additionally, off-target effects of inhibitors against the T3SS, such as destruction of protective members of the microbiota should be rare, given as T3SSs are only found in pathogenic bacteria. Additionally, the development of resistance mutations to cope with the stresses produced by an inhibitor of the T3SS is less likely as such a substance does not affect bacterial viability or structural integrity (12).

A growing number of studies have identified promising inhibitory compounds that target the T3SS of pathogenic bacteria (13–17). While several, such as Aurodox (18), were shown to be selective in inhibiting the T3SS of only one genus, other compounds, including the salicylidene acylhydrazides showed a broader specificity (19–22). Furthermore, some compounds targeted not only the T3SSs but also bacterial motility (23, 24), suggesting that they affect a conserved target in the basal structure shared between virulence-associated T3SSs and the flagellar export system (25).

Here, we used a high throughput screen to identify substances that inhibit T3S-mediated Tir-effector translocation from EPEC into eukaryotic cells. We identified three promising substances which were able to inhibit Tir translocation into host cells without adverse effects on bacterial or eukaryotic cell viability. We were able to show that the inhibitors interfered with intimate bacterial attachment as well as T3SS-dependent cell detachment in response to infection with an EPEC Δ*espZ* mutant. Inhibitors did not affect the amount of LEE-encoded protein expression or motility.

## RESULTS

### Screening of natural and chemical compound libraries identified substances that interfere with effector translocation

Translocation of the EPEC effector protein Tir into the host cell was monitored using an EPEC E2348/69 strain expressing a Tir-β-lactamase (TEM) fusion protein encoded under the native promoter in the EPEC genome (26) and the β-lactamase FRET substrate CCF4-AM, which contains the coumarin- and fluorescein-conjugated β-lactam cephalosporin and is green fluorescent. Using a 96-well plate setup, cells were seeded and subsequently infected with EPEC E2348/69 wildtype or E2346/69 P_LEE5_*tir-blaM.* The bacteria were first grown in T3SS-inducing conditions for two hours and then preincubated in the presence of the screening substances (see **Table 1** for concentrations) for one hour before infection. If translocated into the host cells, the β-lactamase cleaves the CCF4-AM substrate, changing its fluorescence signal from green to blue, which can be quantified in a microtiter plate reader (**Figure 1A**).

**FIG 1.**
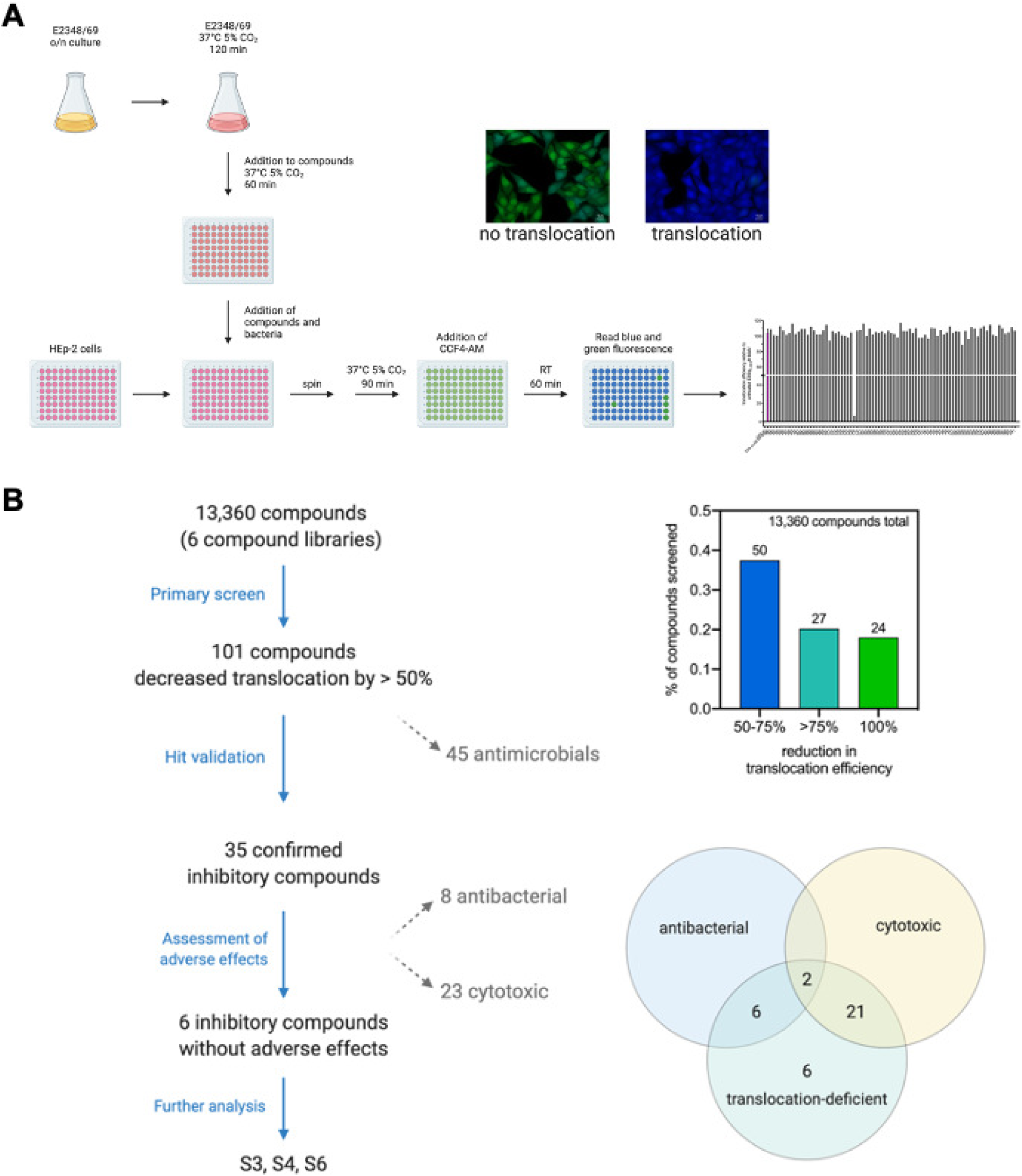
Illustration of inhibitor screen setup and screen summary. For the translocation assay, bacteria were grown overnight in LB broth and diluted 1:50 into DMEM. The infection cultures were grown under T3SS-inducing conditions (37°C, 5% CO_2_) for two hours and subsequently added to 96-well plates containing one µl of the screen sub-stances per well. The bacteria were incubated for an additional hour in the presence of substances before being added to a 96-well plate containing HEp-2 cells. Infection was synchronized by centrifugation. Cells were infected in the presence of inhibitors for 1.5 hours. Bacteria were removed and cells stained with the FRET substrate CCF4-AM for one hour. Translocation efficiency was assessed by measuring blue and green fluorescence (**A**). Of all the libraries screened, compounds that reduced the translocation by more than 50% in the initial screen were further assessed for cytotoxic and antibacterial side-effects. Three inhibitory compounds without adverse effects were used for further analysis (**B**). Figures were created with BioRender.com.

**Table 1:** Screened natural and chemical compound libraries

| Compound library | Source | Number of compounds | Stock solution concentration [mM] | Remarks |
| --- | --- | --- | --- | --- |
| NCh + ExNCI | HIPS, Univ. Tübingen via DZIF | 264 + 352 | 1 and 10 | Microbial secondary metabolites |
| SPECS | Specs, The Netherlands | 2171 | 10 | Chemical synthesis |
| LOPAC | Sigma | 1408 | 10 | Known bioactive compounds |
| Var | Various research groups | 2933 | 5 or 10 | Chemical synthesis |
| EMC | EMC microcollections GmbH, Tübingen | 6232 | 1 | Chemical synthesis |

Using this assay, we screened two natural and four chemical compound libraries with a total of 13,360 substances (**Table 1, S1**). The z’-values ranged between 0.5 and 0.94 with an average z’-value of 0.76 and an average standard to noise ratio (S/N) of 31.8. Of the tested substances, 50 reduced translocation efficiency by 50 to 75%, while 27 were able to reduce the translocation efficiency of Tir-TEM by more than 75% compared to the control (translocation by untreated E2346/69 P_LEE5_*tir-blaM*). 24 compounds showed a reduction of 100% (**Figure 1B, Figure S1, Table S1**). These commonly corresponded to known or published antimicrobial substances including chloramphenicol and rifampicin, thuggacin (27–29), tartrolon (30, 31) as well as derivatives of myxovirescin (32), myxovalargin (33, 34) and sorangicin (35). In total, 45 of the primary hit compounds were identified as antimicrobials or compounds with structural homology to known antibiotics, and were thus excluded from further analyses.

### Effects of primary hit substances on bacterial growth and cell viability

The remaining 56 inhibitory substances identified in the primary translocation screen were then tested for their dose-dependent effect on bacterial growth behavior. For this, overnight cultures of E2348/69 wildtype were diluted to an OD_600_ of 0.02 and grown in the presence of increasing concentrations of each substance (2.5 µM – 50 µM) at 37°C without shaking for eight hours. At regular intervals (every two hours), the optical density of each sample was measured. Of the 56 compounds retested in dose-response assays, only 35 were confirmed to inhibit translocation. Eight of these substances showed an antibacterial effect, reducing the growth of EPEC E2348/69 by >75% at the highest tested concentration (50 µM) (**Figure 1B**).

To further determine whether the inhibitory substances themselves had any effect on the health of the eukaryotic cells (HEp-2), these were seeded in the presence of the substances and incubated at 37°C with 5% CO_2_ for three days. Subsequently, the cell viability was determined by XTT assay. Here, 23 of the tested substances showed marked effects on cellular viability of which two were also shown to have antibiotic activity (**Figure 1B**). In summary, of all tested substances that were neither antimicrobial nor cytotoxic to eukaryotic cells, only six were able to inhibit translocation repeatedly.

### Dose-response assays confirmed three substances that inhibit effector translocation without cytotoxicity to eukaryotic cells or EPEC

The six substances, which were able to inhibit T3SS-mediated translocation of the Tir-TEM fusion repeatedly, were purchased or acquired from their respective sources to allow for further analyses. Repetition of the growth, cell viability and translocation assays with the freshly prepared substances demonstrated that three substances (S3 and S4 from SPECS and S6 from the Var library) robustly inhibited Tir-TEM translocation in a dose-dependent manner (**Figure 2A**) with no adverse effects on bacterial or eukaryotic cell viability (**Figure S2**), with their ID_50_ values given in **Table 2**.

**FIG 2.**
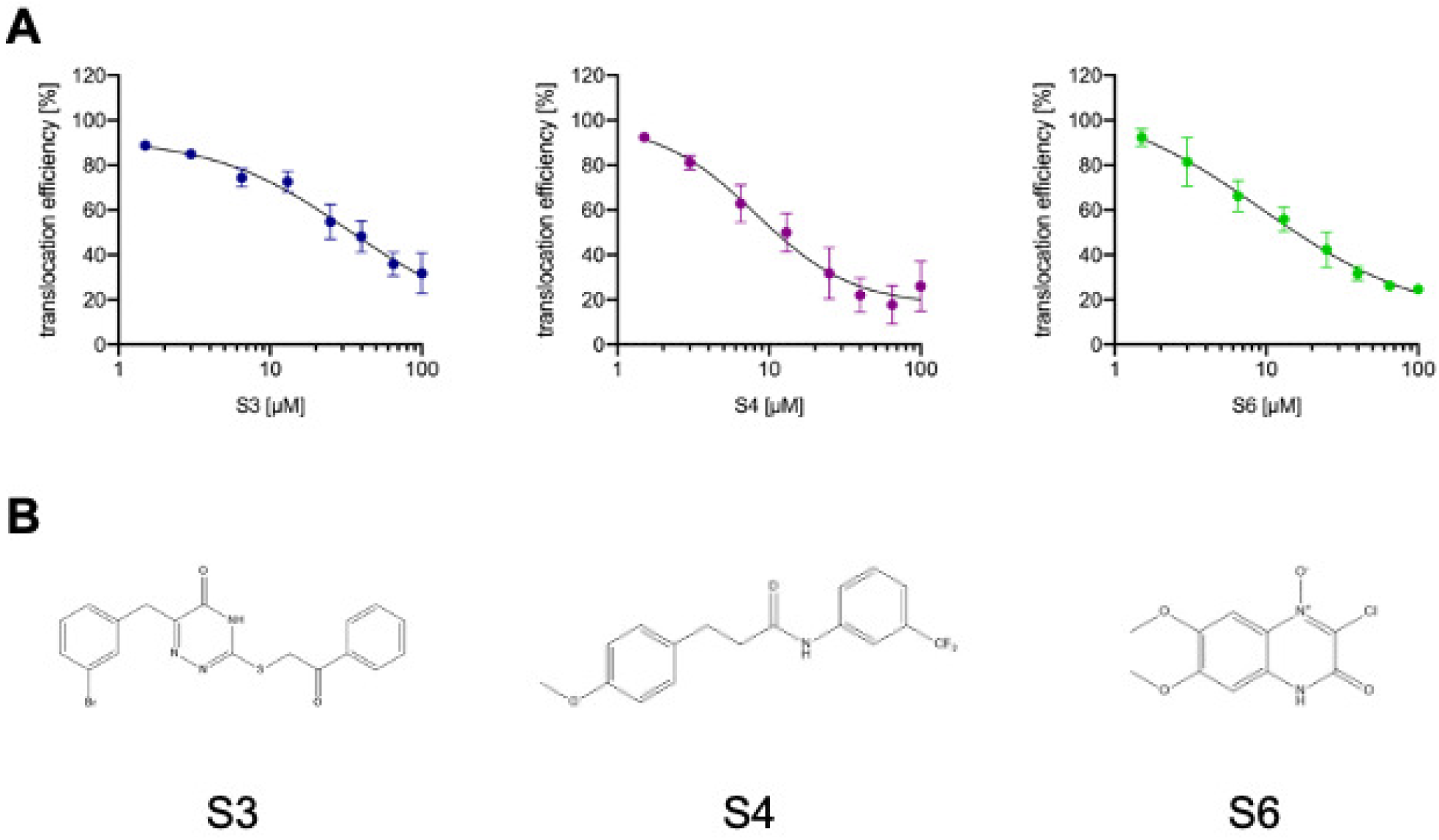
Three substances of the initial high-throughput screen inhibit Tir translocation in a dose-dependent manner. Translocation assays were carried out as described for the high throughput screen. The dose-dependent translocation of Tir-TEM in the presence of the three inhibitory substances identified in the high throughput screen was determined with purchased substances. The ratio of blue/green fluorescence was determined, and values are given as mean (± SEM) of two independent experiments. Curves are analyzed in GraphPad Prism using a non-linear regression fit, variable slope (four parameters) with 95% confidence interval (**A**). Chemical structures of the three substances (drawn using ChemDraw) (**B**).

**Table 2:**
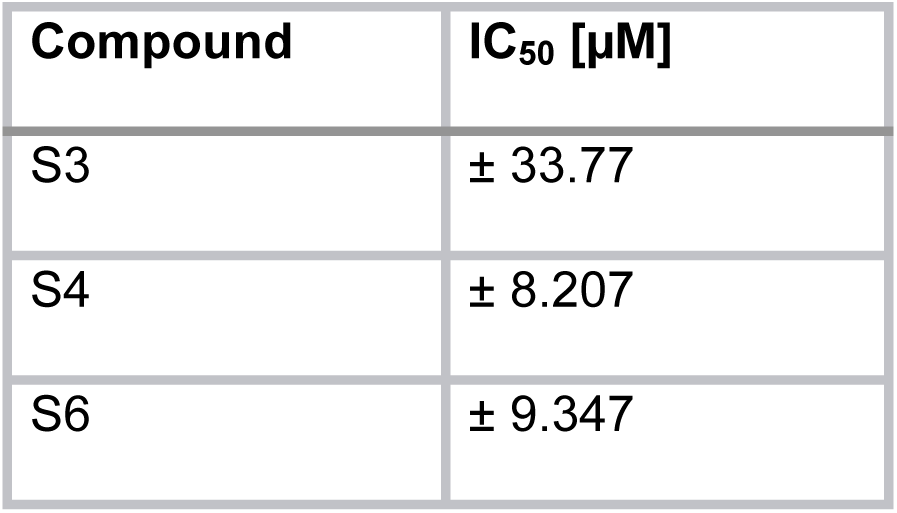
IC_50_ values for translocation efficiency of Tir-TEM as determined with GraphPad Prism using a non-linear regression fit, variable slope (four parameters) with 95% confidence interval

| Compound | IC<sub>50</sub> [μM] |
| --- | --- |
| S3 | ± 33.77 |
| S4 | ± 8.207 |
| S6 | ± 9.347 |

To ensure that the observed inhibition of translocation is not due to an effect on the host cells, preventing host-pathogen interaction, HEp-2 cells were pre-treated with the compounds prior to infection. As shown in **Figure S3**, no inhibition of Tir-TEM translocation was observed.

The chemical structures of the three identified substances are shown in **Figure 2B**. The substances can be categorized as follows: S3 is a 1,2,4-triazine-5-one, S4 is a p-methoxy-hydrocinnamamide, and substance S6 is a 3-chloroquinoxalin-2(*1H*)-one 4-oxide.

### Intimate attachment of EPEC to eukaryotic cells was impaired in the presence of compounds

T3SS-mediated translocation of Tir is followed by its integration into the host cell membrane and interaction with the bacterial outer membrane protein intimin. This induces a phosphorylation cascade that results in the accumulation of F-actin underneath the adherent bacteria and the formation of actin-rich pedestals upon which the bacteria reside (2). HEp-2 cells seeded one day before infection were infected with fluorescently-labelled EPEC strain E2348/69 as described for the translocation assay. Cells were fixed after 1.5 hours of infection, and actin pedestals were visualized by fluorescent actin staining (FAS) using phalloidin. While cells infected with wildtype EPEC in the presence of DMSO (negative control) showed the characteristic actin aggregation where bacteria were attached to the cells, no microcolonies or pedestals could be observed in cells treated with either of the compounds (**Figure 3, Figure S4**). In the presence of the compounds, the majority of the bacteria still visible on the cells were not intimately attached to the surface of the cells, no pedestals were formed (**Figure 3**). No obvious differences were observed between the different compounds. This confirmed previous results showing that all three substances blocked Tir translocation, a prerequisite for pedestal formation.

**FIG 3.**
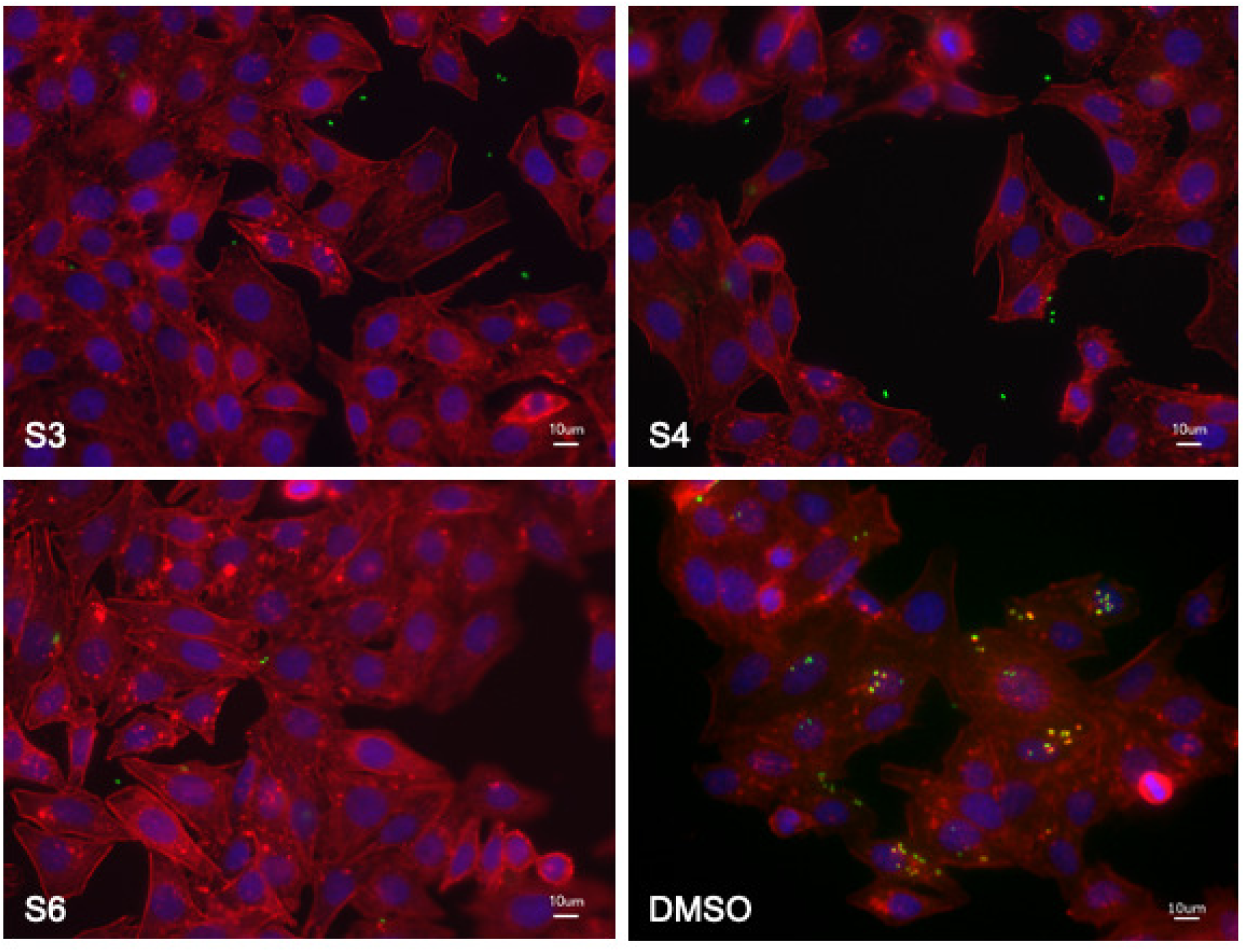
Intimate attachment of EPEC is abolished in the presence of inhibitory substances. HEp-2 cells were infected with EPEC strain E2348/69 pP*_gapdh_amCyan* (green) in the presence of either inhibitory substances or DMSO (control) for 1.5 hours. After rinsing with DPBS, cells were fixed, permeabilized and stained with TRITC-Phalloidin to visualize F-actin (red). Coverslips were mounted with ProLong Diamond mounting medium containing DAPI for staining of DNA (blue). Cells were visualized using a Keyence Biorevo BZ-9000. Scale bars depict 10 µm.

### Substances inhibited translocation-dependent cell detachment

In a previous study, Berger *et al*. identified EspZ as essential for regulating T3SS-dependent protein translocation (36). They showed that a Δ*espZ* mutant induced cell detachment and cell death due to uncontrolled high effector translocation into host cells (36). Here, we used infection of HeLa cells with an EPEC E2348/69 Δ*espZ* mutant in the presence of inhibitors or DMSO to assess whether the inhibitors could block uncontrolled translocation of effector proteins into the host cell and thus inhibit the described cell detachment phenotype. While infection of HeLa cells with wildtype EPEC or the T3SS mutant Δ*escN* had little effect on cell loss, only approximately 15% of cells remained attached after infection with the Δ*espZ* mutant in the presence of DMSO (**Figure 4A and B**). Interestingly, while cell detachment was inhibited in the presence of inhibitors 50 µM S4 or 25 µM S6, which had an average of ∼98% and ∼108% of adherent cells compared to uninfected, 50 µM S3 had a less significant effect on cell loss compared to DMSO-treated cells, with ∼75% of cells remaining after infection (**Figure 4A and B, Figure S5**).

**FIG 4.**
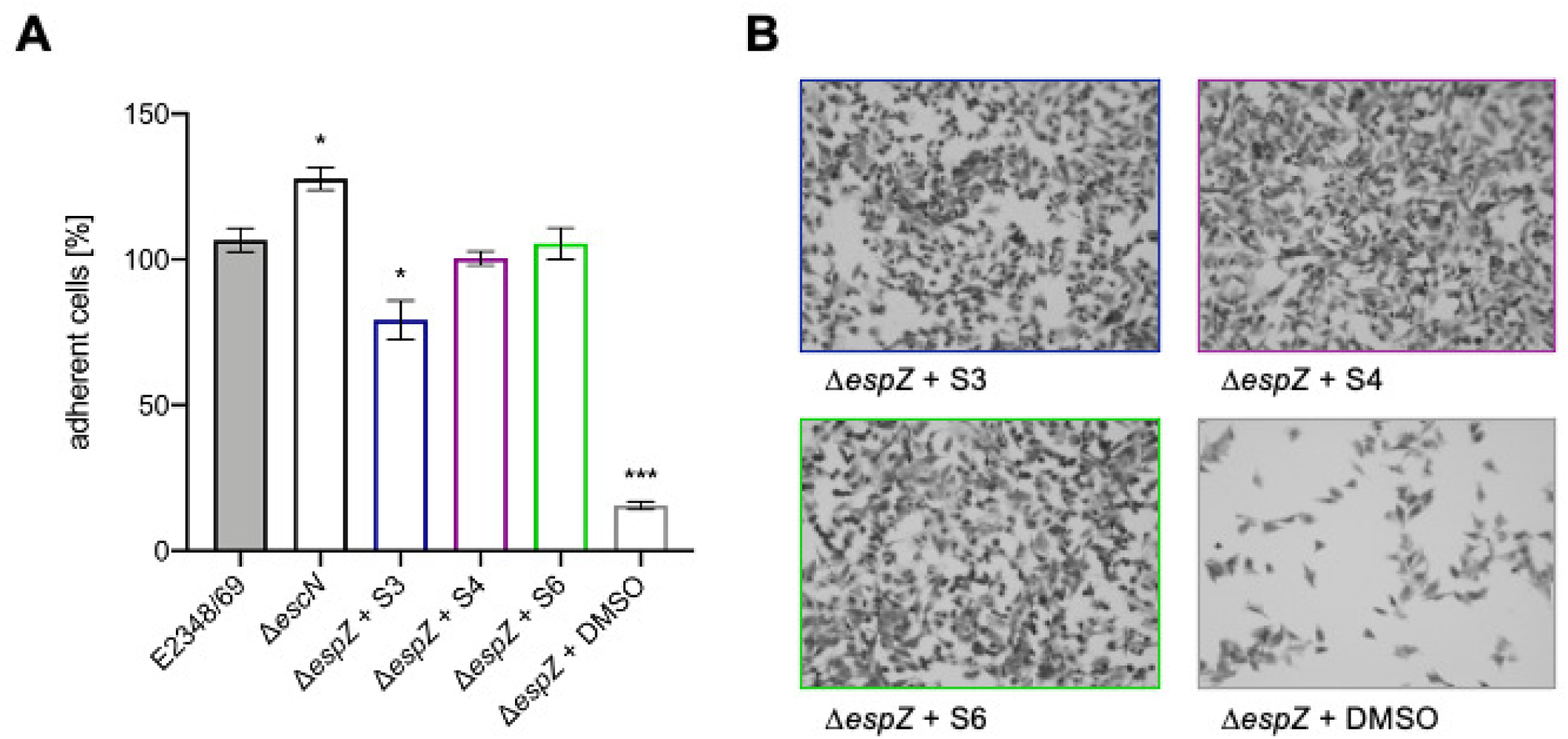
Substances inhibit effector-mediated cell loss. Cells were infected with an EPEC E2348/69 Δ*espZ* mutant in the presence of either 50 µM S3 or S4 and 25 µM S6 or an equivalent amount of DMSO for three hours. Cells were thoroughly washed with DPBS five times every hour. The number of adherent cells after infection was determined by trypsinizing remaining cells at the end of the experiment and counting them in a hemocytometer. The number of cells in uninfected, untreated samples was set to 100%. Shown are the mean values (±SEM) of adherent cells determined in three independent experiments performed at least in triplicate. Statistical analysis was performed via Welch’s t-test (*, P < 0.05; **, P < 0.01; ***, P < 0.001; ****, P < 0.0001), with all treated groups compared to DMSO-treated, Δ*espZ*-infected control cells (**A**). To visualize the amount of cells remaining after three hours, cells were fixed and stained with Hematoxylin Eosin staining solution. Cells were visualized using a Keyence Biorevo BZ-9000 (**B**). Pictures are representative of three independent experiments conducted in triplicate.

### Expression of LEE-encoded proteins remained unaffected in the presence of the inhibitory compounds

The observed effect of the inhibitors on T3SS-mediated translocation of effector proteins such as Tir may be due to a direct effect on the function of the T3S machinery or an indirect inhibition of the process. An indirect effect could reduce the amount of LEE-proteins (either structural components of the T3SS system or translocated effector proteins) expressed in the presence of the substances. To determine whether such an effect on protein expression could be detected, bacteria were grown under T3S-inducing conditions in the presence of the inhibitors, and bacterial cell lysates were assessed by immunoblotting. As can be seen in **Figure 5**, the expression levels of the structural components EspA, EspB and EspD as well as the effector protein Tir (fused to β-lactamase as described above) were comparable in inhibitor-treated bacteria and the DMSO-treated negative control (-) (**Figure 5**). This indicated that either the assembly or the export-translocation function of the effectors of the T3S machinery is inhibited by the compounds.

**FIG 5.**
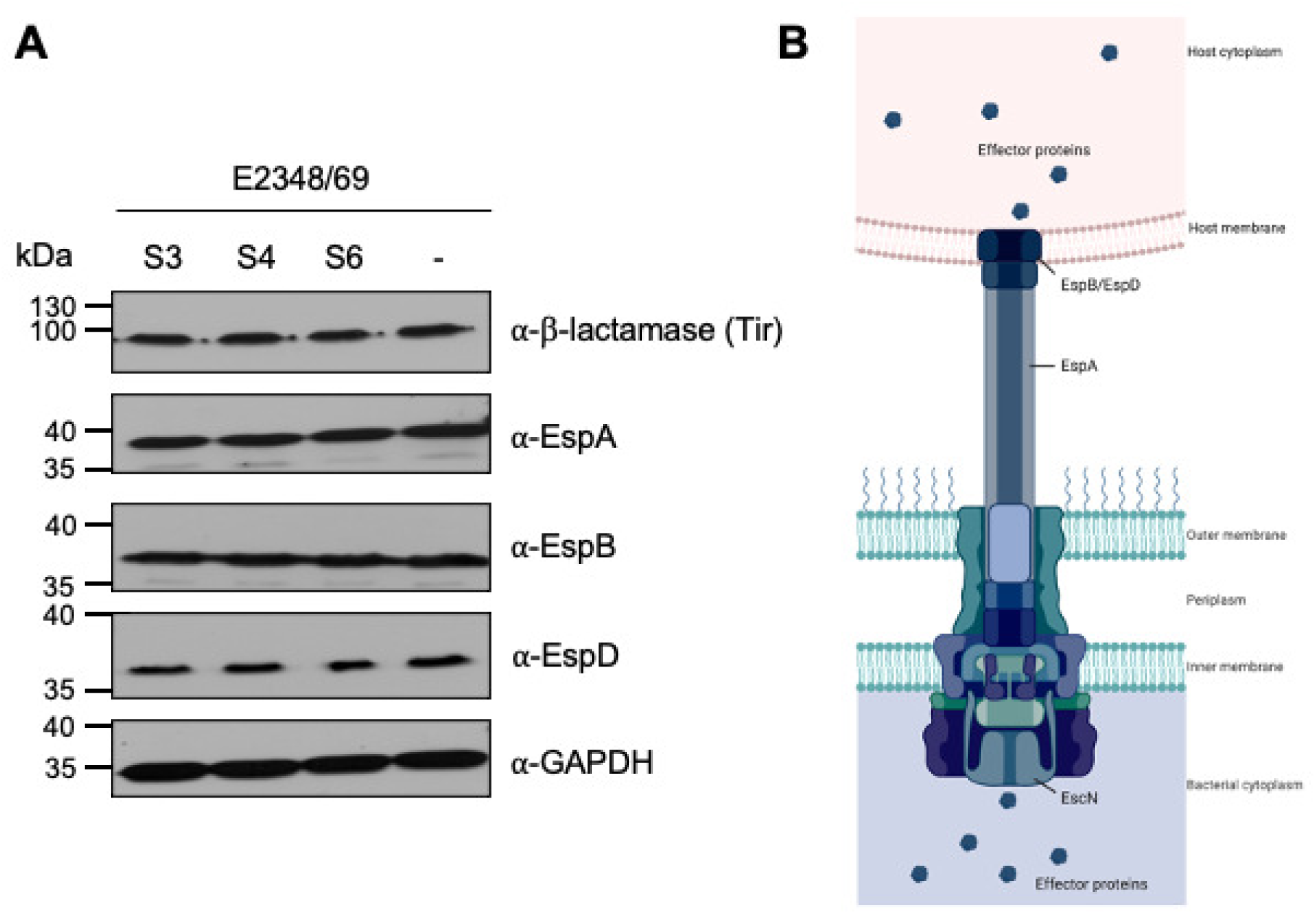
The expression levels of LEE-encoded T3SS proteins are unaffected by the translocation inhibitors. Cell lysates of bacteria grown under T3SS-inducing conditions (37°C, 5% CO_2_) for three hours and equalized for cell numbers were separated by SDS-PAGE. The protein expression levels of the proteins EspA, EspB and EspD, which make up the T3SS filament and translocon pore, as well as the level of the Tir-TEM fusion protein were evaluated by immunoblotting. Depicted blots are representative of at least three independent experiments (**A**). The schematic of the T3SS shows the localization of the analyzed components in the T3SS complex. The figure was created with BioRender.com (**B**).

### The hemolytic activity of EPEC is affected differently by the different inhibitors

T3SS-dependent hemolysis of sheep red blood cells (RBCs) was described for EPEC (37) and has been attributed to the formation of the T3SS translocon pore in the erythrocyte membrane (8). In a previous study, Kimura *et al.* used this phenotype in their high-throughput screen, which identified Aurodox (18). Here, RBCs were infected with EPEC wildtype in the presence or absence of inhibitors and hemolysis was determined after two hours. A Δ*escN* mutant was used as a control for the T3SS-mediated effect, uninfected RBCs were used as a negative, and saponin-treated cells as a positive control. S3 had no effect on hemolysis, while S4 and S6 significant inhibited hemolysis by ∼20% and ∼10%, respectively, compared to DMSO-treated cells (**Figure 6**). This indicated that S3 does not seem to interfere with the assembly and translocon pore formation by the T3SS system.

**FIG 6.**
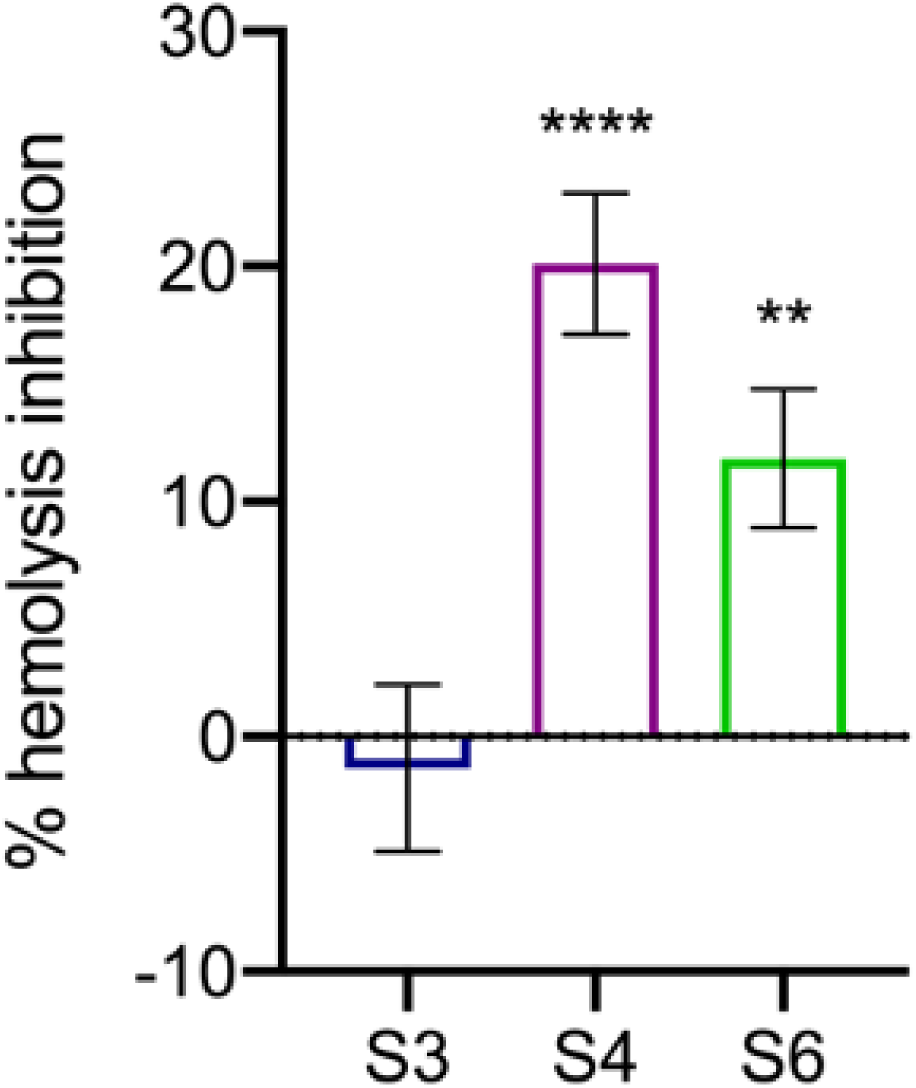
Hemolysis of sheep red blood cells (RBC) was reduced in the presence of S4 and S6 but not by S3. A 3% RBC suspension was incubated with 1×10^8^ bacteria in the presence of either inhibitors or DMSO for two hours. Cells were pelleted, supernatants transferred to 96-well plates, and the amount of hemoglobin released from the cells was determined by measuring absorbance at 543 nm in a plate reader. Shown are the mean values (±SEM) of the percentage of hemolysis inhibition determined in three independent experiments performed in triplicate with all treated groups compared to DMSO-treated, wildtype-infected control cells. Statistical analysis was performed via Welch’s t-test (*, P < 0.05; **, P < 0.01; ***, P < 0.001; ****, P < 0.0001).

### Compounds showed no effect on bacterial motility

Due to the high homology between the basal bodies of the T3SS and flagella (25), a possible effect of the identified compounds on bacterial motility was investigated. Bacterial motility assays were conducted using LB and DMEM soft agar plates supplemented with 50 µM of each substance. As the motility diameter indicates (**Figure 7**), bacteria were more motile on LB plates (∼15 mm) compared to on DMEM plates (∼4 mm). Nevertheless, none of the investigated compounds affected bacterial motility when compared to the negative control (DMSO; **Figure 7**). Bacterial growth in both LB and DMEM was confirmed to be unaffected under the respective conditions (**Figure S6**).

**FIG 7.**
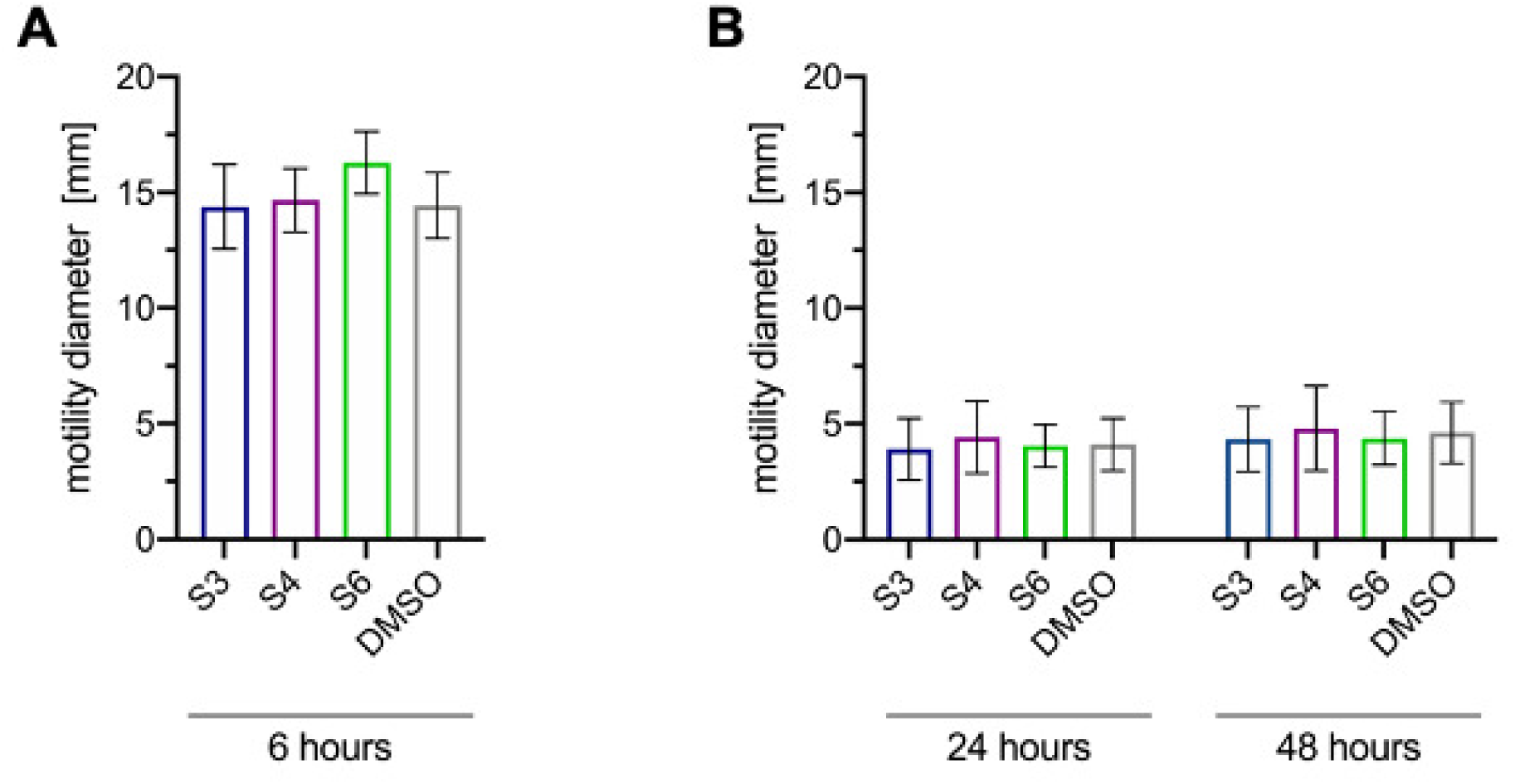
The T3SS inhibiting compounds S3, S4 and S6 do not affect the flagellar T3SS. Bacterial motility was assessed by stab-inoculating bacteria into LB- or DMEM-motility agar containing inhibitors (50 µM S3 or S4, 25 µM S6) or DMSO. After six hours (LB; **A**), 24 hours and 48 hours (DMEM; **B**) the diameters of the motility halo were measured. All assays were carried out three times in triplicate. Differences in diameter in the presence of substances compared to DMSO were not significant. Statistical analysis was performed with Welch’s t-test.

### Inhibitory substances did not induce Shiga toxin production in reporter-gene assays

Previous results of this study suggest that S3, S4 and S6 are effective compounds suppressing the translocation of crucial virulence factors by the T3SS expressed by EPEC but also by EHEC. This could make them to promising candidates for the treatment or the prevention of bloody diarrhea and the development of hemolytic-uremic syndrome (HUS) in infected patients (38, 39), which is associated with the expression of the Shiga toxin (Stx) by EHEC. As many antimicrobial compounds, that induce DNA damage also induce the expression of the Shiga toxin genes (*stx*) which are encoded on a lambdoid prophage, we wanted to ensure that the induction of toxin expression in response to the identified inhibitors can be excluded. To test this, Shiga toxin expression was assessed using *Gaussia* luciferase reporter gene assays in both, *E. coli* K12 (C600) and *Citrobacter rodentium* (DBS100) strains encoding the *Gaussia* luciferase gene (*Gluc*) under the control of the respective *stx2* promoters (40). Ciprofloxacin, a potent inducer of Shiga toxin expression, was used as a positive control. At their tested maximum concentrations, none of the tested inhibitors induced reporter-gene expression in *E. coli* K-12 ϕ*stx2_a_::Gluc* (**Figure 8A**) or *C. rodentium* ϕ*stx2_dact_::Gluc* (**Figure 8B**) compared to DMSO treated control samples. As expected, reporter strains treated with ciprofloxacin showed strong induction of luciferase expression (**Figure 8A and B**). Therefore, it can be concluded that neither of the inhibitors affects Shiga toxin expression, making them safe not only for the treatment of EPEC infections but also as potential drugs against infections with EHEC.

**FIG 8.**
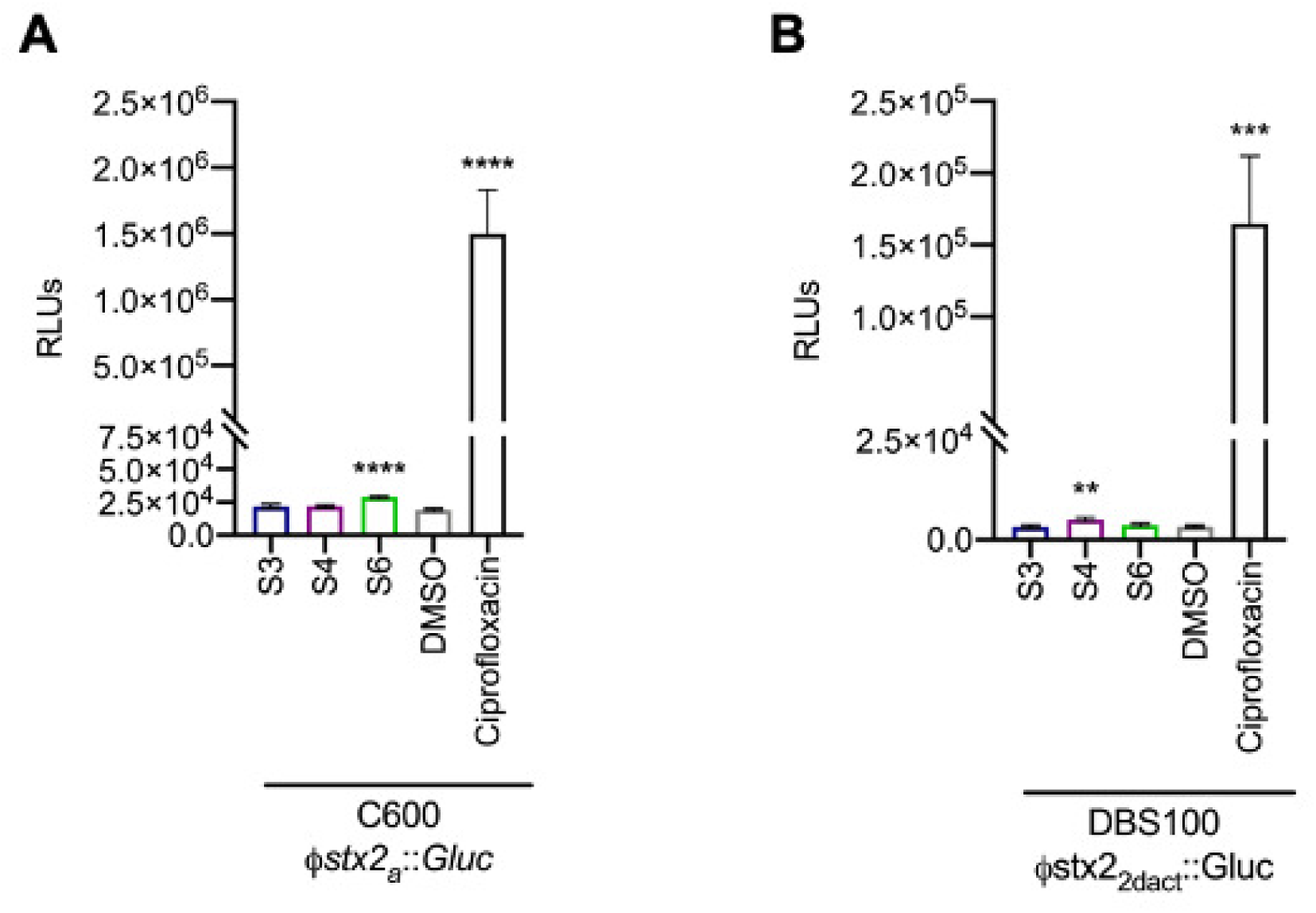
The expression of Shiga toxin is unaffected by the inhibitory compounds. *E. coli* (**A**) or *C. rodentium* (**B**) expressing the *Gaussia* luciferase gene (*Gluc*) under the control of the Shiga toxin promoter were grown overnight in the presence of either compounds, DMSO, or Ciprofloxacin (positive control), equalized and the amount of luciferase released into the supernatant was determined by *Gaussia* luciferase assay. Given are the mean RLU values (±SEM) of three independent experiments performed in triplicate. Statistical analysis was performed via Welch’s t-test (*, P < 0.05; **, P < 0.01; ***, P < 0.001; ****, P < 0.0001), with all treated groups compared to DMSO-treated control.

## DISCUSSION

The dramatic increase in antibiotic resistance in recent years has led to a shift in anti-bacterial strategies towards more focused antivirulence-targeted approaches. Interference with virulence mechanisms instead of bacterial survival is promising to reduce the necessity for resistance development as bacteria will only be impaired in their colonization or pathogenicity but able to survive. With no effect on cell survival, the development of resistance mutations or the necessity for uptake of resistance genes is greatly reduced. Anti-virulence strategies have targeted different virulence factors and virulence-associated processes including motility, adhesion and invasion, effector secretion and inter-bacterial signaling (quorum sensing) (12, 16, 17, 22, 41, 42).

Bacterial secretion systems, especially type III and IV secretion systems, are strongly associated with bacterial virulence, as they are used to inject bacterial effectors into the host cells to modulate disease. The loss of functional secretion systems has, therefore, been associated with a loss of pathogenicity of the respective bacteria, rendering pathogens virtually harmless (11). The phenotypes that are associated with functional T3SSs are diverse. Pathogenic *E. coli* translocate several effector proteins involved in the intimate attachment of the bacteria to the cell surface to withstand being washed off during diarrhea. Other effector proteins mediate the loss of cell-cell junction integrity to invade lower tissues. Furthermore, several effector proteins are involved in the inhibition of innate immune signaling pathways, resulting in the inhibition of cytokine and chemokine release which, in turn, interferes with the recruitment of and clearance by immune cells (43, 44). The formation of the T3SS has also been associated with the formation of pores in and subsequent hemolysis of erythrocytes (8, 37). Moreover, aberrant translocation due to the loss of effectors which control the amount of effector translocation will lead to cell death and detachment (36).

Here, we monitored effector translocation assays in a primary screen to identify small molecule inhibitors that interfere with this process. A Tir-β-lactamase (TEM)-expressing EPEC strain has been used before to study the dynamics of effector translocation in this pathogen (26). Furthermore, this TEM-based assay is commonly used to either screen or confirm identified inhibitory compounds not only in enteropathogenic *E. coli* but also in other T3SS or T4SS-containing pathogens. It has been used to identify or characterize secretion system inhibitors of *Salmonella* (SipB, (23)), *Pseudomonas aeruginosa* (ExoS, (45)) and *Yersinia* (YopE, (46); YopB, (47)), but has also been employed for the T4SSs of *Helicobacter pylori* (48) and *Coxiella burnetii* (49). On the other hand, T3SS-mediated hemolysis was employed as a primary screen in the study, which initially identified the LEE-inhibitor Aurodox (18), while subversion of the host NF-κB inflammatory signaling pathway has been assessed in a study which aimed to identify inhibitors of the *Yersinia* T3SS and identified the inhibitory substance Piericidin A (50).

In our study setup, bacteria were first grown in T3SS-inducing conditions for two hours before exposing them to the inhibitory substances with which they were incubated for an additional hour. Therefore, the inhibitors that were identified in this screen are unlikely to affect the induction of T3SS-gene expression and injectisome formation as after two hours at T3SS-inducing conditions, the T3SSs have already formed. This hypothesis is supported by the fact that the expression of components of the T3SS as well the Tir-TEM fusion protein is unaffected in the presence of the inhibitors (**Figure 5**). All identified T3SS inhibitors S3, S4 and S6 reduced Tir-TEM translocation, effector-dependent cell detachment and A/E lesion formation to a similar extent/in a comparable manner. Most likely, Tir integration into the host cell membrane is impaired as no translocation into the host cell cytosol could be observed for the Tir-TEM fusion protein (**Figures 2, S2**). Furthermore, the absence of pedestals in the presence of the inhibitors conclusively indicated that no intimate interaction occurred between the bacterial outer membrane protein intimin and its receptor, Tir (**Figures 3** and S4) (9, 10). In contrast, T3SS-mediated hemolysis of erythrocytes was only inhibited by S4 and S6, but not by S3. The inhibitory effect of S3 appears to be at the translocation/effector level and does not seem to interfere with T3SS assembly as this inhibitor was unable to inhibit hemolysis of erythrocytes (**Figure 6**). As hemolysis is a consequence of T3 translocon pore formation in the erythrocyte membrane, which is dependent on the translocon pore components EspB and EspD (8, 37), it is possible that S4 and S6 affect translocation of all effectors (including the translocation of EspB and EspD) or that they affect translocon pore-formation *per se* (**Figure 6**). Interestingly, pre-exposure of HEp-2 cells to these inhibitors did not alter the translocation of Tir-TEM by EPEC (**Figure S3**), suggesting that S3, S4 and S6 do not act by influencing host cells. It is likely, that they interfere with the assembly or the function of the T3S apparatus, and it will be interesting to determine the exact mechanism of action of these compounds in future studies.

All three identified inhibitors do not belong to the previously identified classes of T3SS inhibitors. As opposed to previously described T3SS inhibitors (13–17), the inhibitors identified in this study are more specific against the function of the T3SS than previously identified inhibitors. Earlier screens commonly identified inhibitors that affect either gene expression (Aurodox, (51), cytosporone B (52), benzimidazoles (53), sulfonyl amino benzanilides (54), quinolines (55) and salicylidene anilides (24)) or bacterial metabolism (salicylidene acylhydrazides (16), omeprazole ATPase inhibitors (56–58)). While a selective inhibition of LEE (virulence) gene expression is also desirable, inhibitors that affect bacterial metabolism may also result in rapid resistance formation and may also influence the microbiota composition. An effect on the bacterial respiratory chain, which would decrease the formation of the T3SS and thus translocation of effectors was also ruled out for the compounds S3, S4 and S6 identified in this study, as a short-term treatment of bacteria with these inhibitors did not result in significant ATP release (**Figure S7**).

Considering that the identified inhibitors did not affect the expression of the proteins that make up the T3SS or the translocated effector Tir (**Figure 5**), we tested the effect of the inhibitors on the translocation of other effector proteins into the host cell by making use of known effector deletion mutants and their associated phenotypes. Deletion of the gene for the LEE-encoded effector protein EspZ results in aberrant effector translocation into host cells, resulting in host cell detachment due to cell death (36). Preincubation of bacteria in the presence of the inhibitors efficiently reduced cell detachment compared to the positive control (Δ*espZ* + DMSO), suggesting that effector translocation into the host cells was indeed impaired (**Figures 4** and S5).

The homologies between the basal body of the T3SS and the bacterial flagellar basal body are commonly appreciated (25). While an inhibitory effect on bacterial motility would not be undesirable, no effect of any of the newly described substances could be observed (**Figure 7**). This, too, supports the hypothesis that the inhibitors do not interfere with conserved basal body structures of the T3SS, but rather with the assembly of the injectisome or the translocation of the effector proteins. Also, other compounds have been identified that interfere with T3SS function, e.g. (-)-hopeaphenol which seem to localize on the outer membrane and interfere with the closely related T3S apparatus of *Yersinia pseudotuberculosis*, *Pseudomonas aeruginosa* and *Chlamydia trachomatis* (59), and thiazolidinone derivatives that inhibit T3SS in *S. enterica* serovar Typhimurium, *Pseudomonas syringae* and *Francisella tularensis* most likely by targeting the outer membrane secretin component of the T3SS (60). In addition, a picolinic acid derivative and a symmetric dipropionate were found to be active against effector secretion of *Y. pestis* without cytotoxicity against mammalian cells, and notably, they were also active against the translocated intimin receptor Tir in EPEC (46). As their molecular targets are also still unknown, it would be interesting to investigate whether they inhibit identical T3SS functions as the identified compounds S3, S4 or S6.

The effective blockage of T3SS expressed by EPEC as well as by frequent EHEC strains makes the identified inhibitors promising candidates for therapies against hemorrhagic uremic syndrome, which is not curable by common antibiotics such as ciprofloxacin which induce the Shiga toxin genes encoded on lysogenic phages within the bacterial genome (61, 62). As all three inhibitory substances showed no inducing effect on Shiga toxin reporter-gene expression (**Figure 8**), they most likely do not induce/enhance disease progression to HUS. Furthermore, we discovered that even 5-fold excess concentrations (250 µM) of the inhibitors did not affect bacterial growth in MIC studies (**Figure S8**), providing further promise that resistance development against these inhibitory substances may be slow as little selective pressure is induced.

Taken together, our study revealed three potent small molecule inhibitors blocking T3SS-mediated translocation of crucial virulence factors of EPEC/EHEC. As all three compounds interfere with the infection at later stages, e. g. after the T3SS has formed, they constitute promising drugs for therapeutic treatment of the infection, as T3SS-formation will have already occurred once treatment commences.

## Materials and Methods

### Bacterial strains and cell lines

Bacterial strains (listed in **Table S2**) were grown in Luria Bertani (LB; Carl Roth, Germany) or Müller-Hinton (MH; Sigma Aldrich, Germany) broth or DMEM GlutaMAX (Gibco) as indicated. LB overnight cultures were supplemented with antibiotics tetracycline (30 µg/ml), carbenicillin (100 µg/ml) or kanamycin (50 µg/ml) for selection where required.

Cell lines and their respective growth media are given in **Table S3**. Eukaryotic cells were cultured at 37°C, 5% CO_2_.

### Infection of eukaryotic cells

One day before infection, bacteria were inoculated into LB broth and grown at 37°C and 180 rpm overnight. On the day of infection, overnight cultures were diluted 1:75 in DMEM GlutaMAX (Gibco) and grown statically for three hours at 37°C with 5% CO_2_.

HEp-2 cells were washed twice with Dulbecco’s PBS (DPBS; Sigma) and infected with EPEC grown to an OD_600 nm_ of 0.03 for three hours.

### Small molecule compound libraries

All used two natural and four chemical libraries are given in **Table 1** with concentration of stock solutions and sources.

#### Effector translocation assay

For translocation assays, 2×10^5^ HEp-2 (ATCC CCL-23) cells/ml were seeded into black 96-well plates with transparent bottom (Costar, Germany). Bacterial overnight cultures were diluted 1:50 into an appropriate amount of DMEM with GlutaMAX (Gibco, Germany) and incubated for two hours at 37°C 5% CO_2_. 2.5 mM probenecid were added to the cultures which were diluted 1:2 in DMEM with GlutaMAX (Gibco, Germany). 50 µl of the bacterial suspension were subsequently added to the screening plates which contained 1 µl of compound per well (concentrations are given in **Table 1**). These plates were incubated for an additional hour. HEp-2 cells were washed once with Dulbecco’s PBS (DPBS; Sigma, Germany) and 50 µl DMEM with GlutaMAX was added to each well. The bacteria-inhibitor suspension (50 µl per well) was then added to the cells, the plates were centrifuged at 1,000 x g for one minute (Eppendorf Centrifuge 5810R with a A-2-DWP-AT plate rotor) and incubated for 1.5 hours (37°C, 5% CO_2_). Media was then removed, and infected cells were washed twice with DPBS. DMEM with GlutaMAX supplemented with 1 mM HEPES (Biochrom, Germany) and 1x gentamicin (Sigma) was added to the cells and mixed with LifeBLAzer CCF4-AM staining solution (Invitrogen). The plates were then incubated for one hour at room temperature. Subsequently, the fluorescence was determined in a VarioSkan (Fisher Scientific, Germany) or ClarioStar Plus (BMG Labtech, Germany) plate reader using an excitation wavelength of 405 nm (10 nm bandwidth). Emission was detected with 460 nm (20 nm bandwidth, blue fluorescence) and 530 nm (15 nm bandwidth, green fluorescence) filters. Effector translocation was determined by calculating the ratio of blue to green fluorescence (Em_520 nm_/ Em_460 nm_). The translocation efficiency of untreated, EPEC-infected cells was set to 100% while that of untreated EPEC-infected cells was used as a negative control.

#### Analysis of inhibitors on *in vitro* growth and cell viability

Wildtype EPEC E2348/69 was grown overnight in LB at 37°C and resuspended to an optical density (OD_600nm_) of 0.02. 100 µl were added to each well of a 96-well plate containing appropriate amounts of substance or control. Plates were incubated at 37°C without shaking and the OD_600nm_ was determined every two hours for eight hours in a VarioSkan (Fisher Scientific, Germany) or ClarioStar Plus (BMG Labtech, Germany) plate reader.

To determine the effect of inhibitory substances on the viability of eukaryotic cells, 100 µl of a 2×10^4^ cells/ml solution was used to resuspend the inhibitors of interest, and the resulting solution was transferred to a tissue culture treated 96-well plate. Plates were incubated at 37°C, 5% CO_2_ for three days. Subsequently, media was aspirated, and a solution of XTT/PMS (Cell Proliferation Kit II; Merck, Germany) and culture medium was added to the wells. Cells were then incubated at 37°C 5% CO_2_ for an additional two hours after which cell viability was determined at 475 nm in a VarioSkan (Fisher Scientific, Germany) or ClarioStar Plus (BMG Labtech, Germany) plate reader.

#### Fluorescent actin stain (FAS)

Cells seeded on coverslips at 2×10^5^ cells/ml were infected as described above. After infection, cells were washed twice with DPBS, fixed with 4% paraformaldehyde (Sigma, Germany) for 20 min and permeabilized with 0.1% Triton X-100 (OMNI Life Science, Germany) for five minutes at RT. After washing, cells were stained with TRITC-Phalloidin (Sigma Aldrich, Germany), washed again and mounted using Prolong Diamond mounting medium containing DAPI (Thermo Fisher Scientific, Germany). Actin pedestals on cells were visualized using a Keyence Biorevo BZ-9000 (Keyence, Germany).

#### Cell detachment assay

Cell detachment assays were carried out, as described in (36). In short, HeLa cells were seeded 48 hours prior to infection with EPEC primed in the presence or absence of inhibitors. After one hour of infection, cells were washed five times with DPBS, incubated in the presence of inhibitors for an additional hour and washed again five times with DPBS. Cells were trypsinized and counted in a hemocytometer to determine the percentage of remaining cells. For visualization, HeLa cells grown in 8-well removable cell culture chambers (Sarstedt, Germany) or on coverslips were infected and treated as described above. Following the second washing step, cells were fixed with 4% PFA and subsequently stained using the Hematoxylin-Eosin fast staining kit (Carl Roth, Germany) according to (36).

#### Motility assay

LB and DMEM motility agar plates (0.3% agar) containing inhibitory substances at a concentration of 50 µM/ml were stab-inoculated using 2 µl of an EPEC E2348/69 overnight culture and incubated at 37°C overnight. The diameter of the motility halos was documented after 6 (LB) or 24 (DMEM) hours. Triplicate stabs were done per assay for each inhibitor.

#### Protein expression and extraction

For analysis of effector protein expression, LB overnight cultures of bacteria were diluted 1:50 into DMEM GlutaMAX in the presence or absence of inhibitors. After stationary incubation of the cultures at 37°C, 5% CO_2_ for three hours, equalized amounts of cells were harvested by centrifugation at 8,000 x g for 10 minutes. The supernatants were discarded, and the bacterial pellets were resuspended in BugBuster protein extraction reagent (Merck, Germany). The cell suspensions were incubated on a rotary shaker for 20 minutes. SDS sample buffer was added before the samples were boiled at 95°C for 10 minutes before further analysis.

#### SDS-PAGE and immunoblotting

12% SDS-polyacrylamide gels were used to separate proteins with the BioRad MiniPROTEAN Electrophoresis System (BioRad, Germany) followed by transfer of the proteins onto Immobilon FL PVDF membrane (Millipore, Germany). Antibodies against the T3SS components EspA (1:500), EspB (1:10,000) and EspD (1:10,000) (63) and anti-β-lactamase antibody (Abcam; 1:5,000) against the Tir-β-lactamase fusion protein were used to detect protein by immunoblotting. GAPDH (MA5-15738; 1:5,000; ThermoFisherScientific, Germany) was used as a loading control. Using anti-mouse IgG HRP-linked secondary antibody (Cell Signaling Technology, Germany), proteins were detected with WesternLightning ECL Reagent (Perkin Elmer, Germany) and exposure of the membrane to CL-XPosure film (ThermoFisher Scientific, Germany).

#### Hemolysis assay

EPEC overnight cultures were diluted 1:25 in DMEM high glucose without Phenol Red (Sigma, Germany) and grown for three hours at 37°C, 5% CO_2_ in the presence or absence of inhibitors. Sheep RBCs in Alsever (Fiebig Nährstofftechnik, Germany) were washed three times in PBS and resuspended to 5% (vol/vol) in DMEM high glucose without Phenol Red (Sigma, Germany). Bacterial cultures were equalized to 1×10^8^ in 500 µl and added to 500 µl sheep RBCs (5% vol/vol) in two ml Eppendorf tubes. Uninfected RBCs in DMEM were used as a negative control. Total lysis was achieved by the addition of 0.2% saponin to the culture medium. To synchronize infection and mediate bacterial-cell contact, tubes were centrifuged one minute at 2,500 x *g* (Eppendorf MiniSpin) before incubation at 37°C, 5% CO_2_. After two hours, cells were gently resuspended, followed by centrifugation at 2,500 x g for one minute. 100 µl of each supernatant was transferred to a 96-well plate, and the amount of hemoglobin released was assessed at 543 nm in a ClarioStar Plus plate reader (BMG Labtech, Germany). Hemolysis was calculated as the percentage of hemoglobin released by the DMSO-treated wildtype-infected RBCs.

#### Reporter-gene assays

C600 (*stx2*::*Gluc*) and DBS100 (*stx2_dact_*::*Gluc*). Bacteria were inoculated into LB broth and grown at 37°C, 200 rpm for 16 hours in the presence of either the inhibitors at their highest non-cytotoxic concentration, DMSO (negative control; AppliChem) or 40 ng/ml ciprofloxacin (positive control; Sigma). The optical density of each culture was determined, and a volume corresponding to an OD_600nm_ of one (10^9^ cells) was pelleted at 14,000 x *g* for one minute (Eppendorf MiniSpin). 20 µl of supernatant was mixed with 50 µl BioLux *Gaussia* Luciferase Assay substrate (New England Biolabs, Germany) and the luciferase activity was determined in a ClarioStar Plus plate reader (BMG Labtech, Germany) according to the manufacturer’s recommendations. Values were compared to untreated culture supernatant, and LB medium was used as blank.

## Acknowledgements

We thank Ilan Rosenshine for kindly providing the E2348/69 P_LEE5_*tir-blaM* strain and Martin Koeppel and Bärbel Stecher for the use of C600 ϕ*stx2_a_*::*Gluc* and DBS770 ϕ*stx2_dact_::Gluc*. We also thank Sandra Stengel and Amrei Rolof for technical assistance with the HTS. Petra Dersch, Ursula Bilitewski and Sabrina Mühlen are supported by the Deutsche Zentrum für Infektionsforschung (DZIF TTU-GI, TTU 06.801-2, TTU_06_819_00).

